# *Regeneration Rosetta*: An interactive web application to explore regeneration-associated gene expression and chromatin accessibility

**DOI:** 10.1101/632018

**Authors:** Andrea Rau, Sumona P. Dhara, Ava J. Udvadia, Paul L. Auer

## Abstract

Time-course high-throughput assays of gene expression and enhancer usage in zebrafish provide a valuable characterization of the dynamic mechanisms governing gene regulatory programs during CNS axon regeneration. To facilitate the exploration and functional interpretation of a set of fully-processed data on regeneration-associated temporal transcription networks, we have created an interactive web application called *Regeneration Rosetta*. Using either built-in or user-provided lists of genes in one of dozens of supported organisms, our web application facilitates the (1) visualization of clustered temporal expression trends; (2) identification of proximal and distal regions of accessible chromatin to expedite downstream motif analysis; and (3) description of enriched functional gene ontology categories. By enabling a straightforward interrogation of these rich data without extensive bioinformatic expertise, *Regeneration Rosetta* is broadly useful for both a deep investigation of time-dependent regulation during regeneration in zebrafish and hypothesis generation in other organisms.

## Introduction

Axon degeneration accompanying central nervous system (CNS) injury or disease leads to a permanent loss of function in human patients. This is largely due to an inability of mammals to reinitiate axon growth in adult CNS neurons (Crair and Mason 2016). In contrast to mammals, adult teleost fish can fully regenerate CNS axons that reinnervate appropriate targets, enabling functional recovery from CNS injury (Diekmann *et al.* 2015). Interestingly, fish and mammals share common mechanisms for wiring the nervous system during development, and both are known to downregulate developmental growth and guidance signaling pathways during nervous system maturation (Skene 1989; Erskine and Herrera 2014). Thus, what appears to set fish apart is the ability to re-initiate a sustained program of axon growth in response to CNS injury.

Transcriptional changes have long been correlated with the intrinsic capacity for regenerative axon growth (Smith and Skene 1997; Moore and Goldberg 2011). In order to understand the precise mechanisms governing gene regulatory programs during CNS axon regeneration, Dhara et al. (2019) recently identified the dynamic changes in gene expression and enhancer usage in zebrafish over the full time-course of axon regeneration in CNS neurons that are capable of successful regeneration. Adult zebrafish were subjected to optic nerve crush injury, and regenerating retinas were dissected at various time-points post injury in order to identify the interactions among expressed genes, open chromatin, and transcription factor expression during CNS axon regeneration.

These data on regeneration-associated temporal transcription networks in zebrafish represent a rich source of information with wide potential use and insight for the broader regeneration community. To this end, we provide fully processed data from Dhara et al. (2019) in an interactive web application, *Regeneration Rosetta*, as a means to explore, visualize, and functionally interpret regeneration-associated gene expression and chromatin accessibility. Using either built-in lists of differentially expressed (DE) genes from Dhara et al. (2019) or user-provided gene lists in one of 69 supported organisms from Ensembl (Table 1), our web application facilitates (i) customized visualization of clustered temporal expression trends during optic nerve regeneration; (ii) identification of proximal and distal regions of open chromatin relative to the gene list to expedite downstream motif analysis via the MEME suite (Bailey *et al.* 2009); and (iii) gene ontology (GO) functional enrichment analysis. Similarly, using either built-in lists of differentially accessible chromatin from Dhara et al. (2019) or user-provided genomic coordinates of accessible chromatin, the application identifies proximal and distal genes relative to their position and their corresponding enriched GO categories.

**Table 1.**
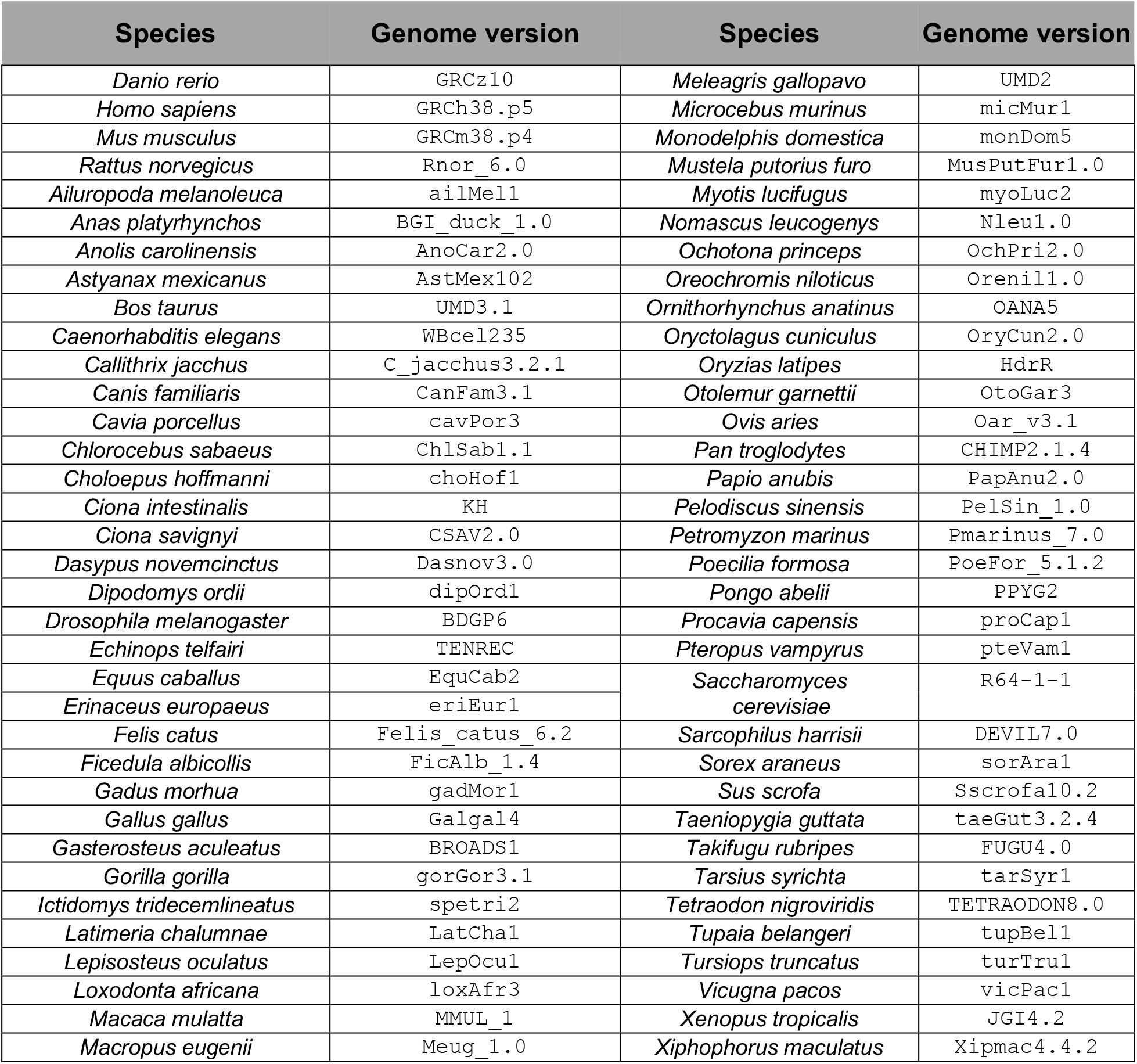
List of supported organisms and their associated genome version for user-provided gene set queries in the *Regeneration Rosetta* app.

The *Regeneration Rosetta* app represents a new paradigm to facilitate data sharing and re-use in the field of regeneration. This type of data sharing directly promotes the National Institutes of Health guidelines for ensuring rigor and reproducibility in pre-clinical research (https://www.nih.gov/research-training/rigor-reproducibility/principles-guidelines-reporting-preclinical-researchas). As large-scale genomic data become more common, tools that allow rapid querying and exploration of fully processed data (without the need for additional coding or pre-processing steps) will be critical in accelerating advances in the regeneration field.

## Materials and Methods

### Experimental design, data generation, and bioinformatic analyses

Comprehensive experimental details may be found in Dhara et al. (2019). Briefly, whole retinas were dissected from 7-9 month old adult zebrafish at 0, 2, 4, 7, or 12 days post injury (dpi), following an optic nerve crush lesion (*n*=3 at each time point). For the RNA-seq data, we tested differential expression with respect to the initial time point (0dpi), controlling the false discovery rate at 5%. For the ATAC-seq data, sequences were aligned (Li and Durbin 2009) to the zebrafish reference and open chromatin regions were called (Zhang *et al.* 2008). For each region with a p-value < 10e-10, a 500bp “peaklet” was defined by anchoring on the mode of the peak signal (Lawrence *et al.* 2013). Chromatin accessibility was quantified by counting the number of overlapping reads for each retained peaklet, and differential accessibility was calculated with respect to the initial time point (0dpi) (Love *et al.* 2014), controlling the false discovery rate at 5%. Software versions and parameters are provided in Dhara et al. (2019). All genomic coordinates and annotations are reported with respect to the GRCz10 Danio rerio genome assembly and Ensembl 90 gene annotation for the zebrafish.

### Integration of RNA-seq and ATAC-seq data

To link regions of accessible chromatin with gene expression, we calculated peaklet-to-gene distance based on the coordinates of the peaklet mode and the gene’s transcription start site (TSS). A proximal peaklet was then defined as one that overlaps the TSS and/or is within ±1kb of the TSS, while a distal peaklet was defined as one within ±100kb of the TSS but not proximal. Users can optionally remove exonic peaklets from these lists, defined as those within 50bp of exonic regions but not overlapping a TSS. To identify genes that are proximal or distal to a given set of accessible chromatin (whether user-provided or through the built-in lists of differentially accessible chromatin provided in the app), users may choose to include all genes or only a subset of those identified to be DE at a particular time point.

For peaklets identified as proximal or distal to the query set of genes, a FASTA file of sequences and BED file of genomic coordinates may be downloaded by the user for further analysis; in addition, a CSV file providing the potentially many-to-many correspondences of proximal and distal peaklets to genes may also be downloaded.

### Gene expression visualization and queries for alternative species

Several built-in gene lists, based on the results described in Dhara et al. (2019), are directly available within the *Regeneration Rosetta* app. These include the lists of DE genes (based on an FDR adjusted p-value < 0.01) compared to 0dpi, as well as pre-identified clusters with expression patterns roughly corresponding to established events in the regeneration process (down-regulation during early-, mid-, or late-regeneration, growth toward the midline, midline crossing, target selection, and brain innervation). Users may also employ the *Regeneration Rosetta* app to explore gene sets for one of 68 species (Table 1) in addition to zebrafish by providing the relevant Ensembl gene IDs (Durinck *et al.* 2009). These converted gene lists can be further narrowed to include only those genes found to be DE post-injury.

For a given set of genes, expression heatmaps using log fold-changes, log transcripts per million (TPM), or Z-scores of these measures are produced using *ComplexHeatmap* (Gu *et al.* 2016), where transcript clusters are identified for a given number of clusters using the K-means algorithm, and rows are ordered within each cluster according to hierarchical clustering (Euclidean distance, complete linkage). Samples may also be hierarchically clustered. A high-resolution heatmap may be resized and downloaded by the user.

### Functional enrichment analysis

The *Regeneration Rosetta* performs on-the-fly functional enrichment analyses of GO terms for Biological Processes (BP), Cellular Components (CC), and Molecular Function (MF) for a given gene set using *topGO* (`weight01` algorithm, Fisher test statistic, and gene universe defined as the set of expressed transcripts from Dhara et al. (2019). P-values are not adjusted for multiple correction, and only GO terms with raw p-values < 0.05 are reported; tables of enriched GO terms are displayed in an HTML table in the app and may be optionally downloaded as a CSV file, Excel spreadsheet, or PDF file. We remark that a list of enriched GO terms must be interpreted with respect to the context of the user-provided gene list. Most users of the *Regeneration Rosetta* would typically make use of lists of regeneration-related genes from one of the supported organisms to understand their behavior in the gene expression and chromatin accessibility data of Dhara et al. (2019); however, if provided with genes unrelated to regeneration, the app will typically return lists of enriched GO terms also unrelated to regeneration. We note that a variety of other ontology databases and enrichment methods exist in the literature, and the interested user could easily export a gene list obtained from the app to other tools, if desired.

### Technical details of the Regeneration Rosetta

The *Regeneration Rosetta* interactive web app was built in R using the *Shiny* and *flexdashboard* packages. In addition to the other software packages already cited above, it makes use of the *data.table* and *RSQLite* R packages for fast data manipulation, *DT* for rendering HTML tables using JavaScript, *readxl* for parsing data from Excel spreadsheets, *dplyr* for data manipulation, and *tokenizers* to convert user-provided gene IDs into tokens. The *Regeneration Rosetta* R/Shiny app is available at http://ls-shiny-prod.uwm.edu/rosetta/. A FAQ page is available directly on the app website. Source code for the *Regeneration Rosetta* is available from GitHub: https://github.com/andreamrau/rosetta. The processed data used within the app are directly located in https://github.com/andreamrau/rosetta/tree/master/data; scripts used to process the raw data from Dhara et al. (2019) may be found at https://github.com/andreamrau/OpticRegen_2019. Archived source code at the time of publication can be found at https://doi.org/10.5281/zenodo.2658771. The *Regeneration Rosetta* app was developed under a GPL-3 software license.

## Results

### Regeneration Rosetta yields insight into cholesterol and lipid biosynthesis regulation during regeneration

Cholesterol biosynthesis pathways were found to be enriched during regeneration in Dhara et al. (2019) and have been previously shown to be important in axon regeneration in mouse (Wang *et al.* 2018). To more deeply investigate their behavior during optic nerve regeneration in zebrafish and demonstrate some of the capabilities of the *Regeneration Rosetta*, we input a list of 125 genes with cholesterol-related GO terms obtained from the Mouse Genome Informatics (MGI) database (Smith *et al.* 2018) (Table S1), corresponding to 232 expressed zebrafish transcripts, into the *Regeneration Rosetta* app (Fig. 1).

**Figure 1:**
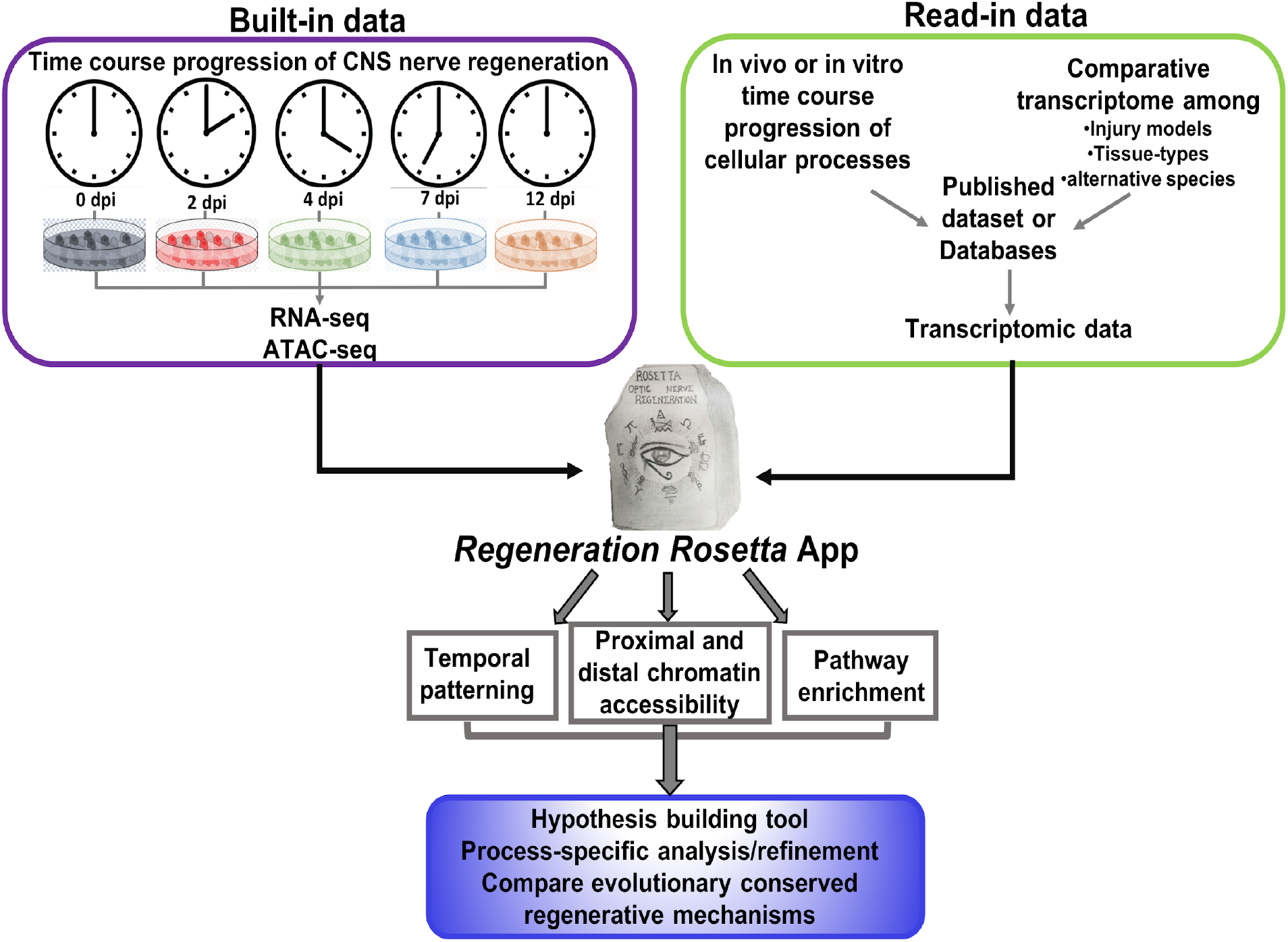
Workflow of *Regeneration Rosetta* app. Workflow for investigating temporal patterning of regeneration-associated genes classified within specific biological processes and/or comparative evolutionary analysis of the conserved mechanism among regenerative species, using the *Regeneration Rosetta* app.

Of these 232 transcripts, 62 were differentially expressed (DE) in Dhara et al. (2019); focusing on this subset of transcripts, the *Regeneration Rosetta* produces a clustered heatmap of expression Z-scores across distinct stages of optic nerve regeneration (Table S2; Figure 2A). Using on the associated GO terms from MGI, we found that the twenty transcripts with peak expression early in regeneration during the phase of growth towards the midline (2-4 dpi) were predominantly enriched in cholesterol metabolic genes, while the majority of those peaking during the midline crossing and target selection phases (4-7 dpi) were enriched in cholesterol biosynthetic pathways (Figure 2B). Interestingly, transcripts differentially down-regulated during regeneration were enriched in negative regulation of cholesterol biosynthetic processes.

**Figure 2:**
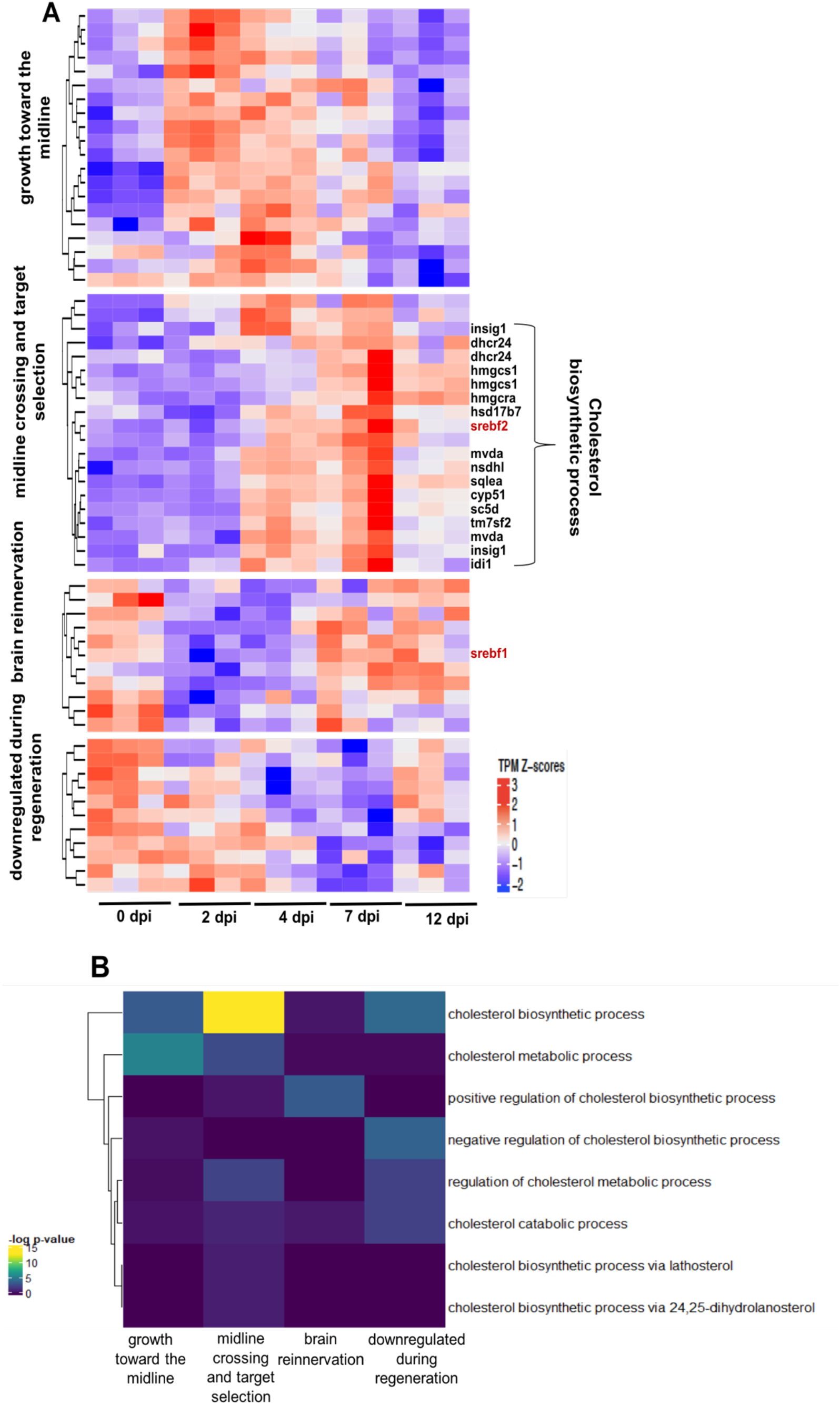
*Regeneration Rosetta* app identifies process-specific analysis after optic nerve injury. (A) Temporal transcript profiles of genes in the cholesterol metabolic pathway. Relative transcript counts from retinas dissected 2-, 4-, 7- and 12-days post injury (dpi) were compared with those from uninjured animals (0 dpi). Transcript expression is presented as TPM Z-scores; putative SREBF2 target genes are indicated to the right of the heatmap (biosynthetic enzymes in black; transcription factors are in red). (B) Specific enrichment of cholesterol metabolic and biosynthetic genes early in regeneration. Fisher’s exact test of over-representation was used to identify cholesterol-related GO-terms correlated with specific stages of regeneration.

Among the cholesterol biosynthetic genes, we observed upregulation during mid-regeneration (4-7 dpi) of SREBF2, a known cholesterol master-regulatory transcription factor (Madison 2016; Smith *et al.* 2018). After downloading the FASTA files of the sequences for peaklets proximal and distal to the 62 DE cholesterol metabolic genes from the *Regeneration Rosetta*, AME motif analysis (McLeay and Bailey 2010) revealed that mostly proximal open chromatin were enriched in the SREBF2 motif resulting in 23% sequence enrichment. We found a number of genes with temporal expression profiles similar to SREBF2 that are proximal to these accessible binding sites, including *dhcr24*, *hmgcs1*, *hsd17b7*, *insig1*, *sqlea*, and *idi1* (Figure 2A). Interestingly, the proximal and distal peaks of *srebf1*, which exhibited peak expression during brain reinnervation (7-12 dpi), were also enriched for SREBF2 motifs; SREBF1 is a transcription factor related to SREBF2 that is known to regulate genes involved in fatty acid synthesis and lipid homeostasis (Ferré and Foufelle 2007; McLeay and Bailey 2010; Ye and DeBose-Boyd 2011), both of which have been shown to be important for axon growth and myelination during neurogenesis (Salles *et al.* 2003; Dietschy and Turley 2004; Ferré and Foufelle 2007). This suggests that SREBF2 could potentially regulate the transcriptional activity of SREBF1, which, in turn, promotes the expression of genes associated with later regenerative processes.

### Regeneration Rosetta provides insight into evolutionary conserved regenerative mechanisms

Nervous system function is dependent on the development of highly specific connections between neurons and their direct targets. The molecular mechanisms regulating this network of connections are highly conserved across evolution (Kaprielian *et al.* 2000; Jiang and Nardelli 2016). Unlike mammals, vertebrates such as fish and amphibians exhibit regenerative abilities of complex tissues and structures. Therefore, comparing regenerative capabilities across species will enable researchers to identify genes and transcriptional networks that are critical to the regenerative program.

To facilitate a cross-species comparison, the *Regeneration Rosetta* app enables queries of patterns of gene expression across injury models with different regenerative capacities. To illustrate, we compared CNS regeneration in lamprey and zebrafish. Following (Kaprielian *et al.* 2000; Herman *et al.* 2018), we used the regenerating lamprey transcriptional profiles from cell bodies located in the brain and spinal cord following spinal cord injury over a course of 12 weeks. This study identified 3,664 and 3,998 differentially expressed regeneration-associated genes at one or more post-injury time points in lamprey brain and spinal cord, respectively. After removing duplicates, we filtered these lists to 2,325 (brain) and 2,519 (spinal cord) differentially expressed genes. We looked for an overlap of genes that were differentially regulated after injury to the zebrafish optic nerve and the lamprey brain and spinal cord. We found 3,712 transcripts (corresponding to 2,150 genes) and 4,076 transcripts (corresponding to 2,404 genes) in the zebrafish optic regeneration data corresponding to the lamprey brain and spinal cord, respectively. After subsetting to those that were differential in the zebrafish (FDR <5%, Wald tests vs 0dpi), we identified 734 (brain) and 846 (spinal cord) differentially expressed genes, 483 were shared between the three data sets (Fig. 3). This suggests a considerable overlap between responses to nerve injury in two different species and three different tissue types.

**Figure 3.**
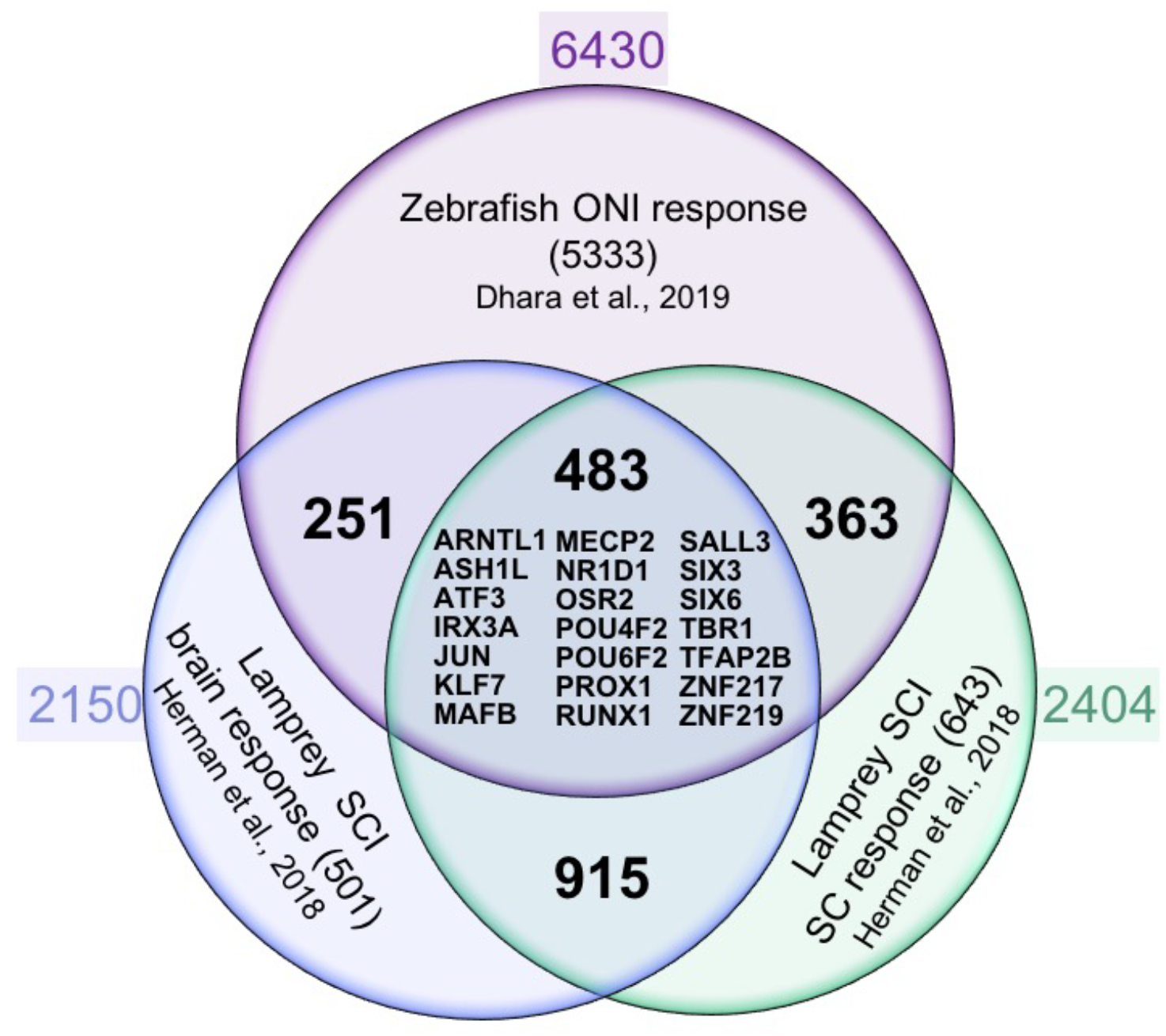
*Regeneration Rosetta* app identifies conserved core regulators of CNS axon regeneration. Venn diagram of axon growth-associated genes from regenerating CNS neurons after zebrafish optic nerve injury (ONI; retina response) and lamprey spinal cord injury (SCI; spinal cord (SC) and brain responses). Approximately 10-20% of regeneration-associated genes are shared between neurons regenerating axons in brain, spinal cord and optic nerve, including a core set of 21 regeneration-associated transcription factor encoding genes that are homologous to human genes (listed in the middle with HGNC gene symbol)

To identify the core transcription factors that could potentially regulate stage-specific regeneration-associated gene transcription in both injury models, we cross referenced the lamprey list of DE transcripts to a recently compiled list of human transcription factors (Lambert *et al.* 2018). We identified 105 brain- and 131 spinal-transcription factor encoding genes that were differentially expressed at one or more post-injury time points (Table S3 and S4). The *Regeneration Rosetta* revealed 32 (brain) and 53 (spinal cord) DE transcription factor encoding genes that were differential in the zebrafish optic nerve regeneration data after subjecting to FDR 5%, Wald tests vs 0dpi. Assessing the combined list of DE transcription factors, we found that 25 transcripts corresponding to 21 transcription factor encoding genes are shared among CNS neuron responses to axonal injury in the lamprey spinal cord and brain, and zebrafish optic nerve (Fig. 3). Thus, the *Regeneration Rosetta* highlights potential regulatory factors driving regeneration-associated gene expression among regeneration capable organisms.

## Discussion

The *Regeneration Rosetta* interactive web app represents a rich resource of fully processed, analyzed, and queryable data from a unique study of regeneration-associated gene expression and chromatin accessibility during optic nerve regeneration in Dhara et al. (2019). The app was a crucial component for generating and interpreting results in Dhara et al. (2019), as it facilitated a deep interrogation of the data that would have otherwise only been possible with extensive bioinformatic expertise. In addition, we have illustrated the broad utility of the *Regeneration Rosetta* app through examples focusing on time-dependent regulation during regeneration for specific biological processes of interest and regenerative mechanisms that are evolutionarily conserved across species and tissue types. All of the source code for running our analyses and implementing the app is posted on GitHub; in this way researchers can modify the code for their own applications or run a local version of the app if desired. As such, the *Regeneration Rosetta* app (and its open-source code) provide a useful a framework for sharing results and data. The *Regeneration Rosetta* app will be widely useful, both for further investigation and interpretation of the data from Dhara et al. (2019) and for hypothesis generation in other organisms. Indeed, we have provided example user-provided gene lists from a variety of published regeneration studies into the app. The lists include regeneration-associated genes from 10 different cell/tissue types from five different species. Our applications allows users to explore how the expression of these genes may change over the course of optic nerve regeneration. We expect these use cases of the app to inform the design of future functional studies that are crucial for translating basic biological insights into new therapeutics for optic nerve injury.

## Acknowledgments

We thank the <u>UWM High Performance Computing (HPC) Service</u> for computing resources used in this work. We are grateful to Maria Replogle for comments on the manuscript. We wish to thank the University of Wisconsin Biotechnology Center DNA Sequencing Facility, Madison for providing RNA and ATAC sample sequencing facilities. We gratefully acknowledge support from the Research Growth Initiative Grant -RGI (to P.L.A. and A.J.U.), University of Wisconsin-Milwaukee. A.R. was supported by the AgreenSkills+ fellowship program, which received funding from the EU’s Seventh Framework Program under grant agreement number FP7-609398 (AgreenSkills+ contract). S.P.D. was supported by the Clifford Mortimer Distinguished Scholar Award, University of Wisconsin-Milwaukee.

