## Supplementary Materials for "*Regeneration Rosetta*: An interactive web application to explore regeneration-associated gene expression and chromatin accessibility"

1. **Table S1**. List of cholesterol metabolic genes from MGI database
2. **Table S2**. List of differentially expressed transcripts during optic nerve regeneration in zebrafish using the MGI cholesterol metabolic gene queries in the Regeneration Rosetta app
3. **Table S3**. List of transcription factor encoding genes from brain following spinal cord injury in lamprey over a course of 12 weeks
4. **Table S4**. List of transcription factor encoding genes from spinal cord following spinal cord injury in lamprey over a course of 12 weeks

| Ensembl ID | MGI Gene ID | Symbol | Name | GO term |
|---|---|---|---|---|
| ENSMUSG00000015243 | MGI:99607 | Abca1 | ATP-binding cassette, sub-family A (ABC1), member 1 | cholesterol metabolic process |
| ENSMUSG00000026944 | MGI:99606 | Abca2 | ATP-binding cassette, sub-family A (ABC1), member 2 | cholesterol metabolic process |
| ENSMUSG00000024030 | MGI:107704 | Abcg1 | ATP binding cassette subfamily G member 1 | cholesterol metabolic process |
| ENSMUSG00000026003 | MGI:87866 | Acadl | acyl-Coenzyme A dehydrogenase, long-chain | regulation of cholesterol metabolic process |
| ENSMUSG00000018574 | MGI:895149 | Acadvl | acyl-Coenzyme A dehydrogenase, very long chain | regulation of cholesterol metabolic process |
| ENSMUSG00000038641 | MGI:2384785 | Akr1d1 | aldo-keto reductase family 1, member D1 | cholesterol catabolic process |
| ENSMUSG00000028553 | MGI:1353627 | Angptl3 | angiopoietin-like 3 | cholesterol metabolic process |
| ENSMUSG00000031996 | MGI:88047 | Aplp2 | amyloid beta (A4) precursor-like protein 2 | cholesterol metabolic process |
| ENSMUSG00000032083 | MGI:88049 | Apoa1 | apolipoprotein A-I | cholesterol biosynthetic process |
| ENSMUSG00000005681 | MGI:88050 | Apoa2 | apolipoprotein A-II | cholesterol metabolic process |
| ENSMUSG00000032080 | MGI:88051 | Apoa4 | apolipoprotein A-IV | cholesterol biosynthetic process |
| ENSMUSG00000032079 | MGI:1913363 | Apoa5 | apolipoprotein A-V | cholesterol biosynthetic process |
| ENSMUSG00000020609 | MGI:88052 | Apob | apolipoprotein B | cholesterol metabolic process |
| ENSMUSG00000042759 | MGI:2176230 | Apobr | apolipoprotein B receptor | cholesterol metabolic process |
| ENSMUSG00000040564 | MGI:88053 | Apoc1 | apolipoprotein C-I | cholesterol metabolic process |
| ENSMUSG00000032081 | MGI:88055 | Apoc3 | apolipoprotein C-III | cholesterol metabolic process |
| ENSMUSG00000002985 | MGI:88057 | Apoe | apolipoprotein E | cholesterol biosynthetic process |
| ENSMUSG00000047631 | MGI:104539 | Apof | apolipoprotein F | cholesterol metabolic process |
| ENSMUSG00000022892 | MGI:88059 | App | amyloid beta (A4) precursor protein | cholesterol metabolic process |
| ENSMUSG00000031982 | MGI:1916115 | Arv1 | ARV1 homolog, fatty acid homeostasis modulator | cholesterol metabolic process |
| ENSMUSG00000027187 | MGI:88271 | Cat | catalase | cholesterol metabolic process |
| ENSMUSG00000034957 | MGI:99480 | Cebpa | CCAAT/enhancer binding protein (C/EBP), alpha | cholesterol metabolic process |
| ENSMUSG00000026818 | MGI:88374 | Cel | carboxyl ester lipase | cholesterol catabolic process |
| ENSMUSG00000056973 | MGI:2148202 | Ces1d | carboxylesterase 1D | cholesterol biosynthetic process |
| ENSMUSG00000041301 | MGI:88388 | Cftr | cystic fibrosis transmembrane conductance regulator | cholesterol biosynthetic process |
| ENSMUSG00000050370 | MGI:1333869 | Ch25h | cholesterol 25-hydroxylase | cholesterol metabolic process |
| ENSMUSG00000032245 | MGI:2159324 | Cln6 | ceroid-lipofuscinosis, neuronal 6 | cholesterol metabolic process |
| ENSMUSG00000026317 | MGI:1349447 | Cln8 | ceroid-lipofuscinosis, neuronal 8 | cholesterol metabolic process |
| ENSMUSG00000026726 | MGI:1931256 | Cubn | cubilin (intrinsic factor-cobalamin receptor) | cholesterol metabolic process |
| ENSMUSG00000018042 | MGI:94893 | Cyb5r3 | cytochrome b5 reductase 3 | cholesterol biosynthetic process |
| ENSMUSG00000028240 | MGI:106091 | Cyp7a1 | cytochrome P450, family 7, subfamily a, polypeptide 1 | cholesterol catabolic process |
| ENSMUSG00000039519 | MGI:104978 | Cyp7b1 | cytochrome P450, family 7, subfamily b, polypeptide 1 | cholesterol metabolic process |
| ENSMUSG00000032323 | MGI:88582 | Cyp11a1 | cytochrome P450, family 11, subfamily a, polypeptide 1 | cholesterol metabolic process |
| ENSMUSG00000075604 | MGI:88583 | Cyp11b1 | cytochrome P450, family 11, subfamily b, polypeptide 1 | cholesterol metabolic process |
| ENSMUSG00000022589 | MGI:88584 | Cyp11b2 | cytochrome P450, family 11, subfamily b, polypeptide 2 | cholesterol metabolic process |
| ENSMUSG00000026170 | MGI:88594 | Cyp27a1 | cytochrome P450, family 27, subfamily a, polypeptide 1 | cholesterol catabolic process |
| ENSMUSG00000023963 | MGI:1927096 | Cyp39a1 | cytochrome P450, family 39, subfamily a, polypeptide 1 | cholesterol catabolic process |
| ENSMUSG00000021259 | MGI:1341877 | Cyp46a1 | cytochrome P450, family 46, subfamily a, polypeptide 1 | cholesterol catabolic process |
| ENSMUSG00000001467 | MGI:106040 | Cyp51 | cytochrome P450, family 51 | cholesterol biosynthetic process |
| ENSMUSG00000030747 | MGI:1915050 | Dgat2 | diacylglycerol O-acyltransferase 2 | regulation of cholesterol metabolic process |
| ENSMUSG00000004815 | MGI:102918 | Dgkq | diacylglycerol kinase, theta | regulation of cholesterol metabolic process |
| ENSMUSG00000058454 | MGI:1298378 | Dhcr7 | 7-dehydrocholesterol reductase | cholesterol biosynthetic process |
| ENSMUSG00000034926 | MGI:1922004 | Dhcr24 | 24-dehydrocholesterol reductase | cholesterol biosynthetic process |
| ENSMUSG00000041544 | MGI:2444403 | Disp3 | dispatched RND transporter family member 3 | cholesterol metabolic process |
| ENSMUSG00000031168 | MGI:107822 | Ebp | phenylalkylamine Ca2+ antagonist (emopamil) binding protein | cholesterol biosynthetic process |
| ENSMUSG00000022040 | MGI:99500 | Ephx2 | epoxide hydrolase 2, cytoplasmic | regulation of cholesterol metabolic process |
| ENSMUSG00000025198 | MGI:2387613 | Erlin1 | ER lipid raft associated 1 | cholesterol metabolic process |
| ENSMUSG00000031483 | MGI:2387215 | Erlin2 | ER lipid raft associated 2 | cholesterol metabolic process |
| ENSMUSG00000021273 | MGI:102706 | Fdft1 | farnesyl diphosphate farnesyl transferase 1 | cholesterol biosynthetic process |
| ENSMUSG00000059743 | MGI:104888 | Fdps | farnesyl diphosphate synthetase | cholesterol biosynthetic process |
| ENSMUSG00000032051 | MGI:103224 | Fdx1 | ferredoxin 1 | cholesterol metabolic process |
| ENSMUSG00000018861 | MGI:104724 | Fdxr | ferredoxin reductase | cholesterol metabolic process |
| ENSMUSG00000024588 | MGI:95513 | Fech | ferrochelatase | cholesterol metabolic process |
| ENSMUSG00000036585 | MGI:95515 | Fgf1 | fibroblast growth factor 1 | sitive regulation of cholesterol biosynthetic proce |
| ENSMUSG00000031594 | MGI:102795 | Fgl1 | fibrinogen-like protein 1 | cholesterol metabolic process |
| ENSMUSG00000031400 | MGI:105979 | G6pdx | glucose-6-phosphate dehydrogenase X-linked | cholesterol biosynthetic process |
| ENSMUSG00000028048 | MGI:95665 | Gba | glucosidase, beta, acid | cholesterol metabolic process |
| ENSMUSG00000028467 | MGI:2654325 | Gba2 | glucosidase beta 2 | cholesterol metabolic process |
| ENSMUSG00000021148 | MGI:3704398 | Gm9745 | predicted gene 9745 | cholesterol biosynthetic process |
| ENSMUSG00000023439 | MGI:95785 | Gnb3 | guanine nucleotide binding protein (G protein), beta 3 | regulation of cholesterol metabolic process |
| ENSMUSG00000034088 | MGI:99256 | Hdlbp | high density lipoprotein (HDL) binding protein | cholesterol metabolic process |
| ENSMUSG00000021670 | MGI:96159 | Hmgcr | 3-hydroxy-3-methylglutaryl-Coenzyme A reductase | cholesterol biosynthetic process |
| ENSMUSG00000093930 | MGI:107592 | Hmgcs1 | 3-hydroxy-3-methylglutaryl-Coenzyme A synthase 1 | cholesterol biosynthetic process |
| ENSMUSG00000027875 | MGI:101939 | Hmgcs2 | 3-hydroxy-3-methylglutaryl-Coenzyme A synthase 2 | cholesterol biosynthetic process |
| ENSMUSG00000029556 | MGI:98504 | Hnf1a | HNF1 homeobox A | cholesterol metabolic process |
| ENSMUSG00000042289 | MGI:2141879 | Hsd3b7 | droxy-delta-5-steroid dehydrogenase, 3 beta- and steroid delta-isomerase | cholesterol catabolic process |
| ENSMUSG00000026675 | MGI:1330808 | Hsd17b7 | hydroxysteroid (17-beta) dehydrogenase 7 | cholesterol biosynthetic process |
| ENSMUSG00000033520 | MGI:2444315 | Idi2 | isopentenyl-diphosphate delta isomerase 2 | cholesterol biosynthetic process |
| ENSMUSG00000000869 | MGI:96556 | Il4 | interleukin 4 | cholesterol metabolic process |
| ENSMUSG00000045294 | MGI:1916289 | Insig1 | insulin induced gene 1 | cholesterol biosynthetic process |
| ENSMUSG00000003721 | MGI:1920249 | Insig2 | insulin induced gene 2 | cholesterol biosynthetic process |
| ENSMUSG00000004880 | MGI:2138281 | Lbr | lamin B receptor | cholesterol biosynthetic process |
| ENSMUSG00000035237 | MGI:96755 | Lcat | lecithin cholesterol acyltransferase | cholesterol metabolic process |
| ENSMUSG00000032193 | MGI:96765 | Ldlr | low density lipoprotein receptor | cholesterol metabolic process |
| ENSMUSG00000037295 | MGI:2140175 | Ldlrap1 | low density lipoprotein receptor adaptor protein 1 | cholesterol metabolic process |
| ENSMUSG00000059201 | MGI:104663 | Lep | leptin | cholesterol metabolic process |
| ENSMUSG00000057722 | MGI:104993 | Lepr | leptin receptor | cholesterol metabolic process |
| ENSMUSG00000023022 | MGI:1920992 | Lima1 | LIM domain and actin binding 1 | cholesterol metabolic process |
| ENSMUSG00000032207 | MGI:96216 | Lipc | lipase, hepatic | cholesterol metabolic process |
| ENSMUSG00000003123 | MGI:96790 | Lipe | lipase, hormone sensitive | cholesterol metabolic process |
| ENSMUSG00000002279 | MGI:1923733 | Lmf1 | lipase maturation factor 1 | regulation of cholesterol metabolic process |
| ENSMUSG00000040249 | MGI:96828 | Lrp1 | low density lipoprotein receptor-related protein 1 | cholesterol metabolic process |
| ENSMUSG00000024913 | MGI:1278315 | Lrp5 | low density lipoprotein receptor-related protein 5 | cholesterol metabolic process |
| ENSMUSG00000033105 | MGI:1336155 | Lss | lanosterol synthase | cholesterol biosynthetic process |
| ENSMUSG00000031835 | MGI:1927235 | Mbtps1 | membrane-bound transcription factor peptidase, site 1 | cholesterol metabolic process |
| ENSMUSG00000031760 | MGI:97173 | Mt3 | metallothionein 3 | cholesterol catabolic process |
| ENSMUSG00000028158 | MGI:106926 | Mttp | microsomal triglyceride transfer protein | cholesterol metabolic process |
| ENSMUSG00000006517 | MGI:2179327 | Mvd | mevalonate (diphospho) decarboxylase | cholesterol biosynthetic process |
| ENSMUSG00000041939 | MGI:107624 | Mvk | mevalonate kinase | cholesterol biosynthetic process |
| ENSMUSG00000038615 | MGI:99421 | Nfe2l1 | nuclear factor, erythroid derived 2,-like 1 | cholesterol metabolic process |
| ENSMUSG00000024413 | MGI:1097712 | Npc1 | NPC intracellular cholesterol transporter 1 | cholesterol metabolic process |
| ENSMUSG00000020447 | MGI:2685089 | Npc1l1 | NPC1 like intracellular cholesterol transporter 1 | cholesterol biosynthetic process |
| ENSMUSG00000021242 | MGI:1915213 | Npc2 | NPC intracellular cholesterol transporter 2 | cholesterol metabolic process |
| ENSMUSG00000031349 | MGI:1099438 | Nsdhl | NAD(P) dependent steroid dehydrogenase-like | cholesterol biosynthetic process |
| ENSMUSG00000044254 | MGI:2140260 | Pcsk9 | proprotein convertase subtilisin/kexin type 9 | cholesterol metabolic process |
| ENSMUSG00000020553 | MGI:107375 | Pctp | phosphatidylcholine transfer protein | cholesterol metabolic process |
| ENSMUSG00000040374 | MGI:107486 | Pex2 | peroxisomal biogenesis factor 2 | regulation of cholesterol biosynthetic process |
| ENSMUSG00000018217 | MGI:97631 | Pmp22 | peripheral myelin protein 22 | cholesterol metabolic process |
| ENSMUSG00000027952 | MGI:1915853 | Pmvk | phosphomevalonate kinase | cholesterol biosynthetic process |
| ENSMUSG00000002588 | MGI:103295 | Pon1 | paraoxonase 1 | cholesterol metabolic process |
| ENSMUSG00000005514 | MGI:97744 | Por | P450 (cytochrome) oxidoreductase | sitive regulation of cholesterol biosynthetic proce |
|  | MGI:104573 | Pphc2 | polyunsaturated fat induced postprandial hypercholesterolemia | cholesterol metabolic process |
| ENSMUSG00000050697 | MGI:2145955 | Prkaa1 | protein kinase, AMP-activated, alpha 1 catalytic subunit | cholesterol biosynthetic process |
| ENSMUSG00000028518 | MGI:1336173 | Prkaa2 | protein kinase, AMP-activated, alpha 2 catalytic subunit | cholesterol biosynthetic process |
| ENSMUSG00000074115 | MGI:98221 | Saa1 | serum amyloid A 1 | cholesterol metabolic process |
| ENSMUSG00000032018 | MGI:1353611 | Sc5d | sterol-C5-desaturase | cholesterol biosynthetic process via lathosterol |
| ENSMUSG00000032485 | MGI:2135958 | Scap | SREBF chaperone | cholesterol metabolic process |
| ENSMUSG00000037936 | MGI:893578 | Scarb1 | scavenger receptor class B, member 1 | cholesterol catabolic process |
| ENSMUSG00000028603 | MGI:98254 | Scp2 | sterol carrier protein 2, liver | sitive regulation of cholesterol biosynthetic proce |
| ENSMUSG00000003585 | MGI:1915065 | Sec14l2 | SEC14-like lipid binding 2 | sitive regulation of cholesterol biosynthetic proce |
| ENSMUSG00000041567 | MGI:1915304 | Serpina12 | ysteine) peptidase inhibitor, clade A (alpha-1 antiproteinase, antitrypsin), | regulation of cholesterol metabolic process |
| ENSMUSG00000037049 | MGI:98325 | Smpd1 | sphingomyelin phosphodiesterase 1, acid lysosomal | cholesterol metabolic process |
| ENSMUSG00000026600 | MGI:104665 | Soat1 | sterol O-acyltransferase 1 | cholesterol metabolic process |
| ENSMUSG00000023045 | MGI:1332226 | Soat2 | sterol O-acyltransferase 2 | cholesterol metabolic process |
| ENSMUSG00000022982 | MGI:98351 | Sod1 | superoxide dismutase 1, soluble | gative regulation of cholesterol biosynthetic proce |
| ENSMUSG00000049313 | MGI:1202296 | Sorl1 | sortilin-related receptor, LDLR class A repeats-containing | cholesterol metabolic process |
| ENSMUSG00000022351 | MGI:109296 | Sqle | squalene epoxidase | cholesterol metabolic process |
| ENSMUSG00000020538 | MGI:107606 | Srebf1 | sterol regulatory element binding transcription factor 1 | cholesterol metabolic process |
| ENSMUSG00000022463 | MGI:107585 | Srebf2 | sterol regulatory element binding factor 2 | cholesterol metabolic process |
| ENSMUSG00000003271 | MGI:1926342 | Sult2b1 | sulfotransferase family, cytosolic, 2B, member 1 | cholesterol metabolic process |
| ENSMUSG00000021779 | MGI:98743 | Thrb | thyroid hormone receptor beta | regulation of cholesterol metabolic process |
| ENSMUSG00000024799 | MGI:1920416 | Tm7sf2 | transmembrane 7 superfamily member 2 | cholesterol biosynthetic process |
| ENSMUSG00000026700 | MGI:104511 | Tnfsf4 | tumor necrosis factor (ligand) superfamily, member 4 | cholesterol metabolic process |
| ENSMUSG00000038172 | MGI:1917113 | Ttc39b | tetratricopeptide repeat domain 39B | regulation of cholesterol metabolic process |
| ENSMUSG00000024924 | MGI:98935 | Vldlr | very low density lipoprotein receptor | cholesterol metabolic process |

**Table S1**. List of cholesterol metabolic genes from MGI database and their associated ensembl ID for user-provided gene set queries in the *Regeneration Rosetta* app.

| Zebrafish Enser | Cluster | Zebrafish_gene | LRT_qval | 2dpi-0dpi_qval | 4dpi-0dpi_qval | 7dpi-0dpi_qval | 12dpi-0dpi_qva | 2dpi-0dpi_LFC | 4dpi-0dpi_LFC | 7dpi-0dpi_LFC | 12dpi-0dpi_LFC |
|---|---|---|---|---|---|---|---|---|---|---|---|
| ENSDART00000 | Growth toward the midline | gba | 0.00060684 | 1.11E-05 | 0.055674455 | 0.928325691 | 0.690320273 | 0.473554928 | 0.246228233 | -0.028007284 | -0.118288896 |
| ENSDART00000 | Growth toward the midline | smpd1 | 0.002176351 | 0.000166524 | 0.430723334 | 0.942817089 | 0.699523141 | 0.404585024 | 0.130204031 | 0.022544887 | -0.113079466 |
| ENSDART00000 | Growth toward the midline | npc2 | 0.002939613 | 0.001556765 | 0.384822365 | 0.914551548 | 0.499923782 | 0.729567353 | 0.287798394 | -0.063690935 | -0.310097689 |
| ENSDART00000 | Growth toward the midline | acadl | 0.002156175 | 0.009322914 | 0.034493954 | 0.905158839 | 0.730001037 | 0.315070362 | 0.266918423 | -0.034640916 | -0.111553388 |
| ENSDART00000 | Growth toward the midline | npc2 | 0.000902446 | 7.69E-08 | 0.058427541 | 0.406017406 | 0.994465692 | 1.578129482 | 0.684715762 | 0.389649636 | 0.031719264 |
| ENSDART00000 | Growth toward the midline | abca1a | 0.015574429 | 0.133882254 | 0.458914664 | 0.033282403 | 0.710864259 | 0.214033988 | 0.130340273 | 0.268396538 | -0.115766571 |
| ENSDART00000 | Growth toward the midline | acadvl | 0.015328252 | 0.01692389 | 0.016316201 | 0.298424368 | 0.979874802 | 0.315023187 | 0.311610409 | 0.17259027 | -0.0260318 |
| ENSDART00000 | Growth toward the midline | nfe2l1a | 0.007828891 | 0.017944516 | 0.011533123 | 0.247377553 | 0.845818106 | 0.652859432 | 0.675562602 | 0.38648833 | -0.18398816 |
| ENSDART00000 | Growth toward the midline | NPC2 (1 of mar | 0.001006824 | 4.68E-07 | 0.001644856 | 0.069931219 | 0.99079012 | 3.182180208 | 2.136779184 | 1.408480393 | 0.09501531 |
| ENSDART00000 | Growth toward the midline | abca1b | 0.004394152 | 0.001218945 | 0.089741646 | 0.655177301 | 0.745138148 | 0.962705135 | 0.587027009 | 0.239926413 | -0.279230387 |
| ENSDART00000 | Growth toward the midline | hdlbpa | 0.002972631 | 0.000342284 | 0.017176287 | 0.099987389 | 0.94653875 | 0.44689202 | 0.322223522 | 0.244841165 | -0.050194137 |
| ENSDART00000 | Growth toward the midline | lepr | 0.000171958 | 2.96E-08 | 1.39E-07 | 1.05E-05 | 0.000856073 | 0.762949533 | 0.720319043 | 0.625239528 | 0.526452567 |
| ENSDART00000 | Growth toward the midline | lepr | 0.000168321 | 5.12E-12 | 2.43E-11 | 1.74E-07 | 0.000959999 | 1.173965688 | 1.129708824 | 0.920018098 | 0.6672678 |
| ENSDART00000 | Growth toward the midline | lepa | 0.00013528 | 6.60E-11 | 3.13E-12 | 3.73E-10 | 0.000724181 | 4.559547685 | 4.778234325 | 4.366752593 | 2.765663683 |
| ENSDART00000 | Growth toward the midline | hsd17b7 | 0.043380755 | 0.044142616 | 0.00678376 | 0.087130748 | 0.336803167 | 2.880332402 | 3.535090877 | 2.510243514 | 2.078786507 |
| ENSDART00000 | Growth toward the midline | acadvl | 0.058646754 | 0.025410864 | 0.014154149 | 0.262740666 | 0.773370237 | 0.465985818 | 0.491501242 | 0.281421395 | 0.167335906 |
| ENSDART00000 | Growth toward the midline | fech | 0.018995329 | 0.991120974 | 0.028094033 | 0.66827511 | 0.988897802 | 0.007306354 | 0.381217581 | -0.12547162 | -0.024425403 |
| ENSDART00000 | Growth toward the midline | si:ch211-93f2.1 | 0.040057894 | 0.694031347 | 0.782937688 | 0.396603565 | 0.049871554 | -0.158550393 | 0.117683559 | -0.256320731 | -0.510696237 |
| ENSDART00000 | Growth toward the midline | cat | 0.008672693 | 0.356612354 | 0.003458029 | 0.865218019 | 0.721963852 | 0.249947463 | 0.551180864 | 0.073389354 | -0.18327225 |
| ENSDART00000 | Growth toward the midline | lrp1aa | 0.02739836 | 0.993779498 | 0.970779906 | 0.337822232 | 0.013674414 | 0.006114382 | -0.023577911 | -0.261635578 | -0.555062861 |
| ENSDART00000 | Midline crossing target selectior | lima1a | 0.014437179 | 0.074429082 | 0.000294565 | 0.007243269 | 0.674291587 | 0.350368785 | 0.588496789 | 0.46589541 | 0.178939257 |
| ENSDART00000 | Midline crossing target selectior | nfe2l1b | 0.013323448 | 0.199387901 | 0.001411736 | 0.007386091 | 0.268361164 | 0.688085235 | 1.301360444 | 1.138008541 | 0.729999096 |
| ENSDART00000 | Midline crossing target selectior | insig1 | 0.008603534 | 0.964439289 | 0.001225581 | 0.028894275 | 0.878598548 | -0.03765091 | 0.839125818 | 0.622258944 | 0.165134159 |
| ENSDART00000 | Midline crossing target selectior | dhcr24 | 0.030292962 | 0.059415464 | 0.022855025 | 0.005780851 | 0.0576435 | 2.392110544 | 2.697063662 | 3.119814675 | 2.653468904 |
| ENSDART00000 | Midline crossing target selectior | dhcr24 | 0.024993104 | 0.36552003 | 0.996650965 | 0.048662488 | 0.947964979 | -0.273134055 | -0.003097208 | 0.45273437 | 0.079225052 |
| ENSDART00000 | Midline crossing target selectior | hmgcs1 | 0.000409858 | 0.764108136 | 0.133189833 | 2.92E-09 | 0.000131252 | 0.250779539 | 0.717953967 | 2.106714123 | 1.525845318 |
| ENSDART00000 | Midline crossing target selectior | hmgcs1 | 0.006978095 | 0.937890417 | 0.779964462 | 0.000788936 | 0.08784359 | -0.061825835 | 0.154559892 | 0.893983325 | 0.609306628 |
| ENSDART00000 | Midline crossing target selectior | hmgcra | 0.005285356 | 0.513462124 | 0.883704521 | 0.0276471 | 0.027858225 | -0.174557714 | 0.058201434 | 0.38988663 | 0.431312576 |
| ENSDART00000 | Midline crossing target selectior | hsd17b7 | 0.001028061 | 0.018553036 | 0.193448 | 0.009655684 | 0.821048076 | -0.488876774 | 0.314769091 | 0.517151777 | 0.148788215 |
| ENSDART00000 | Midline crossing target selectior | srebf2 | 0.003411856 | 0.993271251 | 0.04180132 | 3.17E-05 | 0.09439079 | -0.007030288 | 0.441941306 | 0.773565631 | 0.436893785 |
| ENSDART00000 | Midline crossing target selectior | ldlrb | 0.001252059 | 0.990591906 | 0.013899649 | 7.32E-07 | 0.170571211 | -0.013482381 | 0.739364147 | 1.30137615 | 0.56832803 |
| ENSDART00000 | Midline crossing target selectior | mvda | 0.005340573 | 0.910233515 | 0.013059122 | 0.001717471 | 0.364812696 | 0.177821409 | 1.524901044 | 1.830059782 | 0.927790622 |
| ENSDART00000 | Midline crossing target selectior | nsdhl | 0.073090188 | 0.384843025 | 0.02327355 | 0.01839508 | 0.176504925 | 1.591273733 | 3.017393364 | 3.100136405 | 2.442907405 |
| ENSDART00000 | Midline crossing target selectior | sqlea | 0.000140513 | 0.39714233 | 3.20E-07 | 9.24E-12 | 1.23E-06 | 0.510664466 | 1.875874264 | 2.415595993 | 1.866281513 |
| ENSDART00000 | Midline crossing target selectior | cyp51 | 0.0002611 | 0.979247733 | 9.84E-06 | 5.63E-09 | 0.023589747 | 0.018772193 | 0.850105191 | 1.080881624 | 0.554137375 |
| ENSDART00000 | Midline crossing target selectior | sc5d | 0.009204947 | 0.841975729 | 0.036626452 | 0.003204598 | 0.345217784 | -0.12560715 | 0.619273704 | 0.802648489 | 0.437253012 |
| ENSDART00000 | Midline crossing target selectior | tm7sf2 | 0.008621509 | 0.881675472 | 0.007445135 | 0.000381185 | 0.409104478 | 0.1050645 | 0.758038931 | 0.952561873 | 0.4137868 |
| ENSDART00000 | Midline crossing target selectior | mvda | 0.003470173 | 0.986325295 | 0.000831764 | 0.001919902 | 0.29235693 | -0.034972011 | 1.767545823 | 1.673143467 | 0.929274175 |
| ENSDART00000 | Midline crossing target selectior | insig1 | 0.003330818 | 0.835503343 | 0.008946432 | 0.000357987 | 0.608557059 | -0.148803263 | 0.843708665 | 1.082868772 | 0.366540247 |
| ENSDART00000 | Midline crossing target selectior | idi1 | 0.030676359 | 0.764048984 | 0.028249076 | 0.00420079 | 0.715415577 | 0.080932725 | 0.304370095 | 0.372374682 | 0.127047939 |
| ENSDART00000 | Brain reinnervation | sorl1 | 0.038574021 | 0.498388186 | 0.041236995 | 0.649213893 | 0.785459062 | -0.129978292 | -0.268802979 | -0.097227509 | 0.102143372 |
| ENSDART00000 | Brain reinnervation | si:dkey-71h2.2 | 0.004477335 | 0.002244442 | 2.81E-05 | 0.017235395 | 0.559735938 | -0.547517928 | -0.696070326 | -0.446988451 | -0.222080286 |
| ENSDART00000 | Brain reinnervation | gnb3b | 0.084853017 | 0.047590012 | 0.068454739 | 0.58693258 | 0.95234761 | -0.446677088 | -0.413500571 | -0.183093758 | -0.072200588 |
| ENSDART00000 | Brain reinnervation | scarb1 | 0.022037698 | 0.032429793 | 0.253527644 | 0.929493191 | 0.988391815 | -3.646800721 | -2.308559886 | 0.370802474 | 0.243200954 |
| ENSDART00000 | Brain reinnervation | fdft1 | 0.007609288 | 0.008047735 | 0.010309603 | 0.886937095 | 0.832588895 | -0.467410589 | -0.45021858 | 0.057865823 | -0.125768039 |
| ENSDART00000 | Brain reinnervation | srebf1 | 0.004619636 | 0.002979303 | 0.086540633 | 0.641240969 | 0.988391815 | -0.417658309 | -0.274360919 | 0.114636817 | 0.022214824 |
| ENSDART00000 | Brain reinnervation | hmgcra | 0.001487078 | 0.172515058 | 0.990728292 | 0.020916779 | 0.019457872 | -0.282814218 | -0.006298507 | 0.4014174 | 0.446278745 |
| ENSDART00000 | Brain reinnervation | appb | 0.000152384 | 0.000159341 | 0.006239429 | 0.775755973 | 0.277169566 | -0.473539223 | -0.363304525 | 0.075744005 | 0.226987466 |
| ENSDART00000 | Brain reinnervation | soat2 | 0.016354858 | 0.000255322 | 0.132055603 | 0.461666394 | 0.84389986 | -0.618559497 | -0.31486995 | -0.194806138 | -0.125519338 |
| ENSDART00000 | Brain reinnervation | gnb3a | 0.000657061 | 4.75E-09 | 0.001745536 | 0.036362031 | 0.000171819 | -0.611271729 | -0.360057677 | -0.265096652 | -0.439594606 |
| ENSDART00000 | Brain reinnervation | sec14l8 | 0.069422406 | 0.017565987 | 0.035207725 | 0.548529037 | 0.236825006 | -0.359432343 | -0.324487355 | -0.138175947 | -0.2576341 |
| ENSDART00000 | Downregulated during regenerat | cln6b | 0.003337702 | 0.001075392 | 0.116786127 | 3.48E-06 | 0.292758327 | -0.363769389 | -0.208045101 | -0.481848131 | -0.19203643 |
| ENSDART00000 | Downregulated during regenerat | cyp27a7 | 0.044240808 | 0.014660346 | 0.020052881 | 0.006566642 | 0.202181822 | -0.742372496 | -0.705345887 | -0.793489907 | -0.543448957 |
| ENSDART00000 | Downregulated during regenerat | thrb | 0.063522199 | 0.498401525 | 0.011858621 | 0.029905089 | 0.641189377 | -0.205328436 | -0.495702017 | -0.443190893 | -0.211013858 |
| ENSDART00000 | Downregulated during regenerat | thrb | 0.0223553 | 0.324180982 | 0.002664973 | 0.004383897 | 0.716261895 | -0.193382381 | -0.415286369 | -0.400030228 | -0.136855946 |
| ENSDART00000 | Downregulated during regenerat | ldlrap1b | 0.018733933 | 0.973866391 | 0.052104373 | 0.01071703 | 0.980915101 | -0.019793679 | -0.383158495 | -0.468203159 | -0.036502488 |
| ENSDART00000 | Downregulated during regenerat | gnb3b | 0.060534574 | 0.569323344 | 0.133658566 | 0.010230328 | 0.849977196 | -0.185553204 | -0.343974986 | -0.508555347 | -0.13419791 |
| ENSDART00000 | Downregulated during regenerat | sod1 | 0.030292962 | 0.152465986 | 0.23990086 | 0.002035646 | 0.014580321 | -0.248477917 | -0.212122084 | -0.427881094 | -0.390078569 |
| ENSDART00000 | Downregulated during regenerat | pon2 | 0.003003196 | 0.995047727 | 0.98783272 | 0.003415892 | 0.049780891 | 0.005358778 | -0.010211923 | -0.591626082 | -0.490299784 |
| ENSDART00000 | Downregulated during regenerat | apoeb | 0.046312317 | 0.884739777 | 0.268210448 | 0.014771435 | 0.080531125 | -0.08755429 | -0.343001804 | -0.600108968 | -0.53348823 |
| ENSDART00000 | Downregulated during regenerat | erlin2 | 0.045086702 | 0.999179255 | 0.435223795 | 0.007394961 | 0.862611454 | -0.000462383 | -0.183644452 | -0.434826094 | -0.105981059 |
| ENSDART00000 | Downregulated during regenerat | cln6a | 0.026319966 | 0.744548765 | 0.958508113 | 0.033766616 | 0.848896483 | 0.07921086 | -0.018544414 | -0.27320282 | -0.083540486 |

**Table S2**. List of differentially expressed transcripts during optic nerve regeneration in zebrafish using the MGI cholesterol metabolic gene queries in the Regeneration Rosetta app. The result table includes the zebrafish ensembl ID, DE genes based on Only DE (padj<5%, Wald tests vs 0dpi) for each regenerating timepoints 2, 4, 7 , 12 dpi.

| Brain Official Gene Symbol | Lamprey Gene ID | DNA Binding Domain |
|---|---|---|
| TFAP2B | PMZ_0018996-RA | AP-2 |
| AKNA | PMZ_0011406-RA | AT hook |
| ASH1L | PMZ_0005066-RA | AT hook |
| SGSM2 | PMZ_0017332-RA | BED ZF |
| ZBED1 | PMZ_0016473-RA | BED ZF |
| AHR | PMZ_0009169-RA | bHLH |
| ARNTL | PMZ_0022526-RA | bHLH |
| BHLHE40 | PMZ_0015665-RA | bHLH |
| CLOCK | PMZ_0006274-RA | bHLH |
| MLXIP | PMZ_0011574-RA | bHLH |
| MLXIPL | PMZ_0011572-RA | bHLH |
| NCOA3 | PMZ_0005855-RA | bHLH |
| NEUROD1 | PMZ_0005582-RA | bHLH |
| PTF1A | PMZ_0023241-RA | bHLH |
| SCX | PMZ_0011122-RA | bHLH |
| TWIST1 | PMZ_0008383-RA | bHLH |
| TWIST2 | PMZ_0004583-RA | bHLH |
| ATF3 | PMZ_0005968-RA | bZIP |
| CEBPA | PMZ_0019312-RA | bZIP |
| CREM | PMZ_0012457-RA | bZIP |
| FOS | PMZ_0001200-RA | bZIP |
| FOSL2 | PMZ_0023077-RA | bZIP |
| JUN | PMZ_0015901-RA | bZIP |
| MAFB | PMZ_0014543-RA | bZIP |
| NFE2 | PMZ_0014947-RA | bZIP |
| NFIL3 | PMZ_0019135-RA | bZIP |
| DZIP1 | PMZ_0013304-RA | C2H2 ZF |
| EEA1 | PMZ_0012072-RA | C2H2 ZF |
| IKZF5 | PMZ_0001000-RA | C2H2 ZF |
| KLF1 | PMZ_0020730-RA | C2H2 ZF |
| KLF10 | PMZ_0012571-RA | C2H2 ZF |
| KLF15 | PMZ_0002919-RA | C2H2 ZF |
| KLF7 | PMZ_0008885-RA | C2H2 ZF |
| OSR2 | PMZ_0006091-RA | C2H2 ZF |
| PEG3 | PMZ_0002813-RA | C2H2 ZF |
| PRDM1 | PMZ_0020972-RA | C2H2 ZF |
| PRDM5 | PMZ_0005403-RA | C2H2 ZF |
| PRDM8 | PMZ_0005757-RA | C2H2 ZF |
| REST | PMZ_0007778-RA | C2H2 ZF |
| SALL3 | PMZ_0012241-RA | C2H2 ZF |
| ZBTB42 | PMZ_0017837-RA | C2H2 ZF |
| ZNF135 | PMZ_0019928-RA | C2H2 ZF |
| ZNF217 | PMZ_0000126-RA | C2H2 ZF |
| ZNF219 | PMZ_0000147-RA | C2H2 ZF |
| ZNF22 | PMZ_0011057-RA | C2H2 ZF |
| ZNF239 | PMZ_0018206-RA | C2H2 ZF |
| ZNF676 | PMZ_0019476-RA | C2H2 ZF |
| ZNF704 | PMZ_0017942-RA | C2H2 ZF |
| ZNF841 | PMZ_0002784-RA | C2H2 ZF |
| ZEB2 | PMZ_0017801-RA | C2H2 ZF; Homeodomain |
| TRERF1 | PMZ_0003066-RA | C2H2 ZF; Myb/SANT |
| RBPJL | PMZ_0019954-RA | CSL |
| KDM2A | PMZ_0023537-RA | CxxC |
| KDM2B | PMZ_0005926-RA | CxxC |
| EHF | PMZ_0011596-RA | Ets |

| | | |
|---|---|---|
| ETS1 | PMZ_0023551-RA | Ets |
| ETV7 | PMZ_0019267-RA | Ets |
| FOXB2 | PMZ_0010602-RA | Forkhead |
| FOXG1 | PMZ_0000494-RA | Forkhead |
| FOXK1 | PMZ_0012299-RA | Forkhead |
| FOXK2 | PMZ_0009477-RA | Forkhead |
| FOXL2 | PMZ_0010037-RA | Forkhead |
| GATA2 | PMZ_0008156-RA | GATA |
| GCM2 | PMZ_0010350-RA | GCM |
| SOX10 | PMZ_0020526-RA | HMG/Sox |
| SRY | PMZ_0006678-RA | HMG/Sox |
| ADNP | PMZ_0015678-RA | Homeodomain |
| DBX1 | PMZ_0001656-RA | Homeodomain |
| DLX5 | PMZ_0012874-RA | Homeodomain |
| HOXB6 | PMZ_0019528-RA | Homeodomain |
| HOXB8 | PMZ_0019779-RA | Homeodomain |
| HOXC9 | PMZ_0002315-RA | Homeodomain |
| HOXD8 | PMZ_0006066-RA | Homeodomain |
| HOXD9 | PMZ_0001980-RA | Homeodomain |
| IRX3 | PMZ_0019709-RA | Homeodomain |
| LHX1 | PMZ_0018111-RA | Homeodomain |
| MKX | PMZ_0013726-RA | Homeodomain |
| NKX6-1 | PMZ_0024089-RA | Homeodomain |
| OTX2 | PMZ_0000784-RA | Homeodomain |
| PHOX2B | PMZ_0023008-RA | Homeodomain |
| SIX1 | PMZ_0022507-RA | Homeodomain |
| SIX3 | PMZ_0017270-RA | Homeodomain |
| SIX6 | PMZ_0015477-RA | Homeodomain |
| TPRX1 | PMZ_0023099-RA | Homeodomain |
| POU4F2 | PMZ_0005302-RA | Homeodomain; POU |
| POU6F2 | PMZ_0004395-RA | Homeodomain; POU |
| PIN1 | PMZ_0005768-RA | MBD |
| MECP2 | PMZ_0011998-RA | MBD; AT hook |
| ESR1 | PMZ_0002173-RA | Nuclear receptor |
| HNF4A | PMZ_0002236-RA | Nuclear receptor |
| NR1D1 | PMZ_0012318-RA | Nuclear receptor |
| NR1D2 | PMZ_0018839-RA | Nuclear receptor |
| PPARA | PMZ_0011032-RA | Nuclear receptor |
| RORC | PMZ_0011373-RA | Nuclear receptor |
| PROX1 | PMZ_0012095-RA | Prospero |
| NFATC3 | PMZ_0011590-RA | Rel |
| REL | PMZ_0002455-RA | Rel |
| RUNX1 | PMZ_0019080-RA | Runt |
| DEAF1 | PMZ_0007366-RA | SAND |
| NFIA | PMZ_0005871-RA | SMAD |
| TBR1 | PMZ_0005862-RA | T-box |
| TBX20 | PMZ_0017953-RA | T-box |
| DR1 | PMZ_0012187-RA | Unknown |
| LTF | PMZ_0007403-RA | Unknown |
| SON | PMZ_0022239-RA | Unknown |

**Table S3**. List of transcription factor encoding genes from brain following spinal cord injury in lamprey over a course of 12 weeks (derived from Herman et al., 2018). This table includes the official gene symbol corresponding to lamprey gene ID for the transcription factors with known DNA binding domain for user-provided gene set queries in the *Regeneration Rosetta* app.

| Spinal cord Official Gene Symbol | Lamprey Gene ID | DNA binding domain |
|---|---|---|
| TFAP2B | PMZ_0018996-RA | AP-2 |
| ASH1L | PMZ_0005066-RA | AT hook |
| SETBP1 | PMZ_0010700-RA | AT hook |
| ZBED1 | PMZ_0016473-RA | BED ZF |
| ARNTL | PMZ_0022526-RA | bHLH |
| BHLHE41 | PMZ_0006661-RA | bHLH |
| EPAS1 | PMZ_0007089-RA | bHLH |
| MAX | PMZ_0008167-RA | bHLH |
| MITF | PMZ_0022356-RA | bHLH |
| MLXIP | PMZ_0011574-RA | bHLH |
| MLXIPL | PMZ_0011572-RA | bHLH |
| MSC | PMZ_0003667-RA | bHLH |
| MSGN1 | PMZ_0006006-RA | bHLH |
| MYCN | PMZ_0004744-RA | bHLH |
| NCOA3 | PMZ_0005855-RA | bHLH |
| NEUROD1 | PMZ_0005582-RA | bHLH |
| NPAS3 | PMZ_0007275-RA | bHLH |
| TFE3 | PMZ_0015238-RA | bHLH |
| TWIST1 | PMZ_0003348-RA | bHLH |
| TWIST2 | PMZ_0004583-RA | bHLH |
| ATF3 | PMZ_0021004-RA | bZIP |
| ATF7 | PMZ_0023367-RA | bZIP |
| CEBPA | PMZ_0019312-RA | bZIP |
| CREM | PMZ_0012457-RA | bZIP |
| FOS | PMZ_0019104-RA | bZIP |
| FOSL2 | PMZ_0023077-RA | bZIP |
| HLF | PMZ_0020258-RA | bZIP |
| JUN | PMZ_0015901-RA | bZIP |
| MAFB | PMZ_0014543-RA | bZIP |
| NFE2L2 | PMZ_0009505-RA | bZIP |
| EEA1 | PMZ_0012072-RA | C2H2 ZF |
| EGR1 | PMZ_0018081-RA | C2H2 ZF |
| GLI2 | PMZ_0003580-RA | C2H2 ZF |
| GLIS2 | PMZ_0013642-RA | C2H2 ZF |
| IKZF4 | PMZ_0007774-RA | C2H2 ZF |
| KLF1 | PMZ_0020730-RA | C2H2 ZF |
| KLF7 | PMZ_0013729-RA | C2H2 ZF |
| MAZ | PMZ_0009919-RA | C2H2 ZF |
| MYT1L | PMZ_0020295-RA | C2H2 ZF |
| OSR1 | PMZ_0011713-RA | C2H2 ZF |
| OSR2 | PMZ_0006091-RA | C2H2 ZF |
| PRDM8 | PMZ_0005757-RA | C2H2 ZF |
| SALL3 | PMZ_0012241-RA | C2H2 ZF |
| ZBTB16 | PMZ_0004848-RA | C2H2 ZF |
| ZBTB42 | PMZ_0017837-RA | C2H2 ZF |
| ZFAT | PMZ_0006891-RA | C2H2 ZF |
| ZNF16 | PMZ_0012820-RA | C2H2 ZF |
| ZNF217 | PMZ_0000126-RA | C2H2 ZF |
| ZNF219 | PMZ_0000147-RA | C2H2 ZF |
| ZNF22 | PMZ_0011057-RA | C2H2 ZF |
| ZNF267 | PMZ_0016475-RA | C2H2 ZF |
| ZNF367 | PMZ_0019541-RA | C2H2 ZF |
| ZNF385B | PMZ_0005569-RA | C2H2 ZF |
| ZNF385D | PMZ_0008540-RA | C2H2 ZF |
| ZNF596 | PMZ_0019979-RA | C2H2 ZF |
| ZNF676 | PMZ_0019476-RA | C2H2 ZF |
| ZNF704 | PMZ_0019042-RA | C2H2 ZF |
| ZNF710 | PMZ_0002701-RA | C2H2 ZF |
| ZNF841 | PMZ_0002784-RA | C2H2 ZF |
| ZUFSP | PMZ_0014878-RA | C2H2 ZF |
| CAMTA1 | PMZ_0016476-RA | CG-1 |
| KDM2B | PMZ_0005926-RA | CxxC |
| E2F3 | PMZ_0017731-RA | E2F |
| E2F7 | PMZ_0007982-RA | E2F |
| EBF1 | PMZ_0003202-RA | EBF1 |
| EBF2 | PMZ_0003201-RA | EBF1 |
| EHF | PMZ_0011596-RA | Ets |
| ETV6 | PMZ_0019268-RA | Ets |

| | | |
|---|---|---|
| ETV7 | PMZ_0019267-RA | Ets |
| FOXB2 | PMZ_0010602-RA | Forkhead |
| FOXF2 | PMZ_0003163-RA | Forkhead |
| FOXG1 | PMZ_0000494-RA | Forkhead |
| GCM2 | PMZ_0003269-RA | GCM |
| TFCP2 | PMZ_0015026-RA | Grainyhead |
| SOX11 | PMZ_0001911-RA | HMG/Sox |
| ADNP | PMZ_0015678-RA | Homeodomain |
| DLX1 | PMZ_0021876-RA | Homeodomain |
| DLX5 | PMZ_0012874-RA | Homeodomain |
| DMBX1 | PMZ_0015945-RA | Homeodomain |
| EMX1 | PMZ_0000550-RA | Homeodomain |
| HOXB6 | PMZ_0019528-RA | Homeodomain |
| HOXB8 | PMZ_0019779-RA | Homeodomain |
| HOXC9 | PMZ_0002315-RA | Homeodomain |
| HOXD8 | PMZ_0006066-RA | Homeodomain |
| HOXD9 | PMZ_0001980-RA | Homeodomain |
| IRX2 | PMZ_0020478-RA | Homeodomain |
| IRX3 | PMZ_0019709-RA | Homeodomain |
| IRX5 | PMZ_0012177-RA | Homeodomain |
| LHX1 | PMZ_0018111-RA | Homeodomain |
| MEIS2 | PMZ_0020921-RA | Homeodomain |
| MKX | PMZ_0013726-RA | Homeodomain |
| OTX2 | PMZ_0000784-RA | Homeodomain |
| PHOX2B | PMZ_0023008-RA | Homeodomain |
| SIX1 | PMZ_0022507-RA | Homeodomain |
| SIX3 | PMZ_0017270-RA | Homeodomain |
| SIX4 | PMZ_0014005-RA | Homeodomain |
| SIX6 | PMZ_0015477-RA | Homeodomain |
| TPRX1 | PMZ_0023099-RA | Homeodomain |
| POU4F2 | PMZ_0005302-RA | Homeodomain; POU |
| POU6F2 | PMZ_0004393-RA | Homeodomain; POU |
| MEF2A | PMZ_0017273-RA | MADS box |
| MBD2 | PMZ_0013621-RA | MBD |
| PIN1 | PMZ_0005768-RA | MBD |
| MECP2 | PMZ_0011998-RA | MBD; AT hook |
| DMTF1 | PMZ_0014946-RA | Myb/SANT |
| ESR1 | PMZ_0002173-RA | Nuclear receptor |
| ESR2 | PMZ_0001332-RA | Nuclear receptor |
| ESRRG | PMZ_0017464-RA | Nuclear receptor |
| HNF4A | PMZ_0002236-RA | Nuclear receptor |
| NR1D1 | PMZ_0012318-RA | Nuclear receptor |
| NR1H4 | PMZ_0006651-RA | Nuclear receptor |
| NR2F1 | PMZ_0005031-RA | Nuclear receptor |
| PPARA | PMZ_0011032-RA | Nuclear receptor |
| RARA | PMZ_0010334-RA | Nuclear receptor |
| RARB | PMZ_0010333-RA | Nuclear receptor |
| PROX1 | PMZ_0012095-RA | Prospero |
| REL | PMZ_0002455-RA | Rel |
| RUNX1 | PMZ_0019080-RA | Runt |
| RUNX2 | PMZ_0006895-RA | Runt |
| DEAF1 | PMZ_0007365-RA | SAND |
| GMEB2 | PMZ_0000511-RA | SAND |
| NFIA | PMZ_0005872-RA | SMAD |
| STAT4 | PMZ_0013733-RA | STAT |
| TBR1 | PMZ_0005862-RA | T-box |
| TBX18 | PMZ_0022118-RA | T-box |
| TBX20 | PMZ_0009881-RA | T-box |
| DR1 | PMZ_0004286-RA | Unknown |
| LTF | PMZ_0007403-RA | Unknown |
| RBCK1 | PMZ_0019160-RA | Unknown |
| SON | PMZ_0022239-RA | Unknown |
| TSC22D1 | PMZ_0021850-RA | Unknown |

**Table S4**. List of transcription factor encoding genes from spinal cord following spinal cord injury in lamprey over a course of 12 weeks (derived from Herman et al., 2018). This table includes the official gene symbol corresponding to lamprey gene ID for the transcription factors with known DNA binding domain for user-provided gene set queries in the *Regeneration Rosetta* app.
